# Live fast and die young: walleye populations adapt their life cycle to degraded lakes of the Canadian clay belt

**DOI:** 10.1101/2025.08.27.672400

**Authors:** Patrice Blaney, Pascal Sirois, Martin Bélanger, Eva C. Enders, Marta Gabriele, Guillaume Grosbois

## Abstract

Despite its widespread distribution in North America, many populations of walleye (*Sander vitreus*) declined to the point where restoration measures, including restocking, are necessary. In this study, we compared population structure and dietary composition of walleye populations in two degraded lakes and two non-degraded lakes in the Abitibi-Témiscamingue region of Quebec, Canada. Food resources were assessed for walleye larvae, young-of-the-year, juveniles, and adults. Growth and relative abundance were also quantified for young-of-the-year, juvenile, and adult walleye. Young-of-the-year were more abundant and grew faster in degraded lakes compared to non-degraded controls, benefiting from high populations of spring zooplankton, which are a critical larval resource. The simplified food webs in degraded lakes lacked pollution-sensitive macroinvertebrates, which resulted in walleye diets being even more dominated by fish. Although juveniles and adults were equally or more abundant in degraded lakes compared to control lakes, premature adult mortality compromised population stability. We recommend focusing on improving adult fish habitat, manage for prey species, and review fishing regulations to enhance the survival of mature walleye and ensure sustainable populations.

## Introduction

Recreational anglers in Canada catch more walleye (*Sander vitreus*) than any other fish species. Walleye account for 26% of recreational fish caught and 55% of the dollar value of freshwater commercial fisheries (2010–2020 averages) (DFO 2015; 2022). However, the degradation of spawning sites, decreased water quality, introduction of invasive aquatic species, and overfishing of stocks have reduced walleye populations, with negative consequences for commercial and recreational fisheries (Rooney and Paterson 2009; Bozek et al. 2011a). For example, in Wisconsin (USA), natural recruitment of walleye in lakes has declined from 68% in 1993 to 37% in 2018 (Raabe et al. 2020). Between 700 million and 1 billion walleyes are stocked annually in North America (Raabe et al. 2020), but the success of stocking is difficult to predict and varies greatly with ecosystem characteristics (Raabe et al. 2020). Therefore, investments in restocking by fisheries managers do not guarantee success due to the lack of information needed to predict stocking outcomes.

The self-sustainability of a walleye population in a lake depends on several interconnected factors, including interspecific interactions, habitat quality and type, and habitat connectivity (Bozek et al. 2011a; Raabe et al. 2020). The walleye life cycle is divided into five main stages: egg, larvae, young-of-the-year (age 0, or 0 +), juvenile (age 1 until sexual maturation), and adult. Each life stage has specific resource needs (Bozek et al. 2011a), and those needs must be accounted for when assessing the potential of freshwater ecosystems to sustain fish populations (Raabe et al. 2020).

The Canadian Clay Belt, which extends from eastern Manitoba to western Quebec, is a region where walleye is widely distributed and abundant. This region contains many turbid lakes, to which walleyes are particularly well adapted due to the *tapetum lucidum,* a subretinal structure in their eyes that enables them to feed in low-visibility environments (Ryder 1977; Vandenbyllaardt et al. 1991). However, human activities such as mining have led to local population declines, prompting a restocking with larvae or autumnal young-of-the-year (Kerr 2008). Metals released by mining activities may have direct impacts on fish physiology and indirect, food-web-mediated impacts on fish populations (Campbell et al. 2003). For example, reduced availability of key prey such as cladocerans and macroinvertebrates can hinder growth and survival during early life stages (Leis and Fox 1996; Sherwood et al. 2000; Campbell et al. 2003; Bozek et al. 2011b; McDonnell and Roth 2014).

Natural causes account for the majority of mortality in early life stages of walleye. As walleye grow, natural mortality decreases while fishing-related mortality increases (Hansen et al. 2011). Recreational fishing can also slow the recovery of predatory species (Dainys et al. 2022). Therefore, it is crucial to measure the impacts of recreational fishing on the survival of adult walleye, especially in lakes close to urban areas where the fishing pressure is high (Kaemingk et al. 2020). The cumulative impacts of human activities occurring in a region (e.g., urbanization, industrial development, recreational fishing) is expected to affect both habitat quality and the sustainability of fish populations, including walleye.

The **general objective** of this study was to determine whether lakes degraded by human activities in the Canadian Clay Belt offer the necessary ecological conditions to allow the self-sustainability of walleye populations. We hypothesized that non-degraded lakes would offer more favorable ecological conditions for all life stages of walleye when compared with degraded lakes. We further hypothesized that if key ecological functions (such as critical food resources) were to be maintained in degraded lakes, populations would continue to persist. We tested these hypotheses by describing the food resources available to larvae, young-of-the-year, juveniles and adults. Furthermore, we quantified the relative abundance and measured the growth of walleye young-of-the-year, juveniles, and adults.

## Methodology

### Study area

This study was conducted in four lakes located near to the city of Rouyn-Noranda in the Abitibi-Témiscamingue region of Quebec, Canada. Located in the western balsam fir-white birch bioclimatic subdomain, this region has a subpolar, subhumid, and continental climate (Blouin and Berger 2002; Guimond et al. 2024). The average annual temperature is 2.0 °C, and 27% of the 895 mm annual precipitation falls as snow, which represents up to 2.4 m of snowfall. (MELCCFP 2024a). The region’s 20,000 lakes are typically frozen from December to May (Beaulne et al. 2012; Canadian Cryospheric Information Network 2024; Grosbois et al. 2024).

The high density of lakes in Abitibi-Témiscamingue can be attributed to glaciolacustrine deposits left by Lake Barlow-Ojibway following the last ice age (Blouin and Berger 2002). These deposits have formed poorly drained clay soils that promote runoff and lake creation. These lakes are naturally turbid due to the presence of clay particles and tannins from the boreal forest (Grosbois et al. 2023; Hasan et al. 2023).

The region’s lakes have been impacted by mining activities that began in the early 1900s with the discovery of copper and gold deposits (SNQAT 1981). A regional gradient of metal contamination originates in downtown Rouyn-Noranda, where several mines have conducted their operations (Laflamme et al. 2000). Urbanization has also altered the state of the lakes. Sewage discharges polluted aquatic ecosystems until 1978, when the Quebec government began the Wastewater Purification Program (MDDELCC 2014). Several lakes close to Rouyn-Noranda are still degraded, particularly those situated near urban areas. Despite their condition, these lakes continue to provide economic benefits and serve as sources of drinking water. Moreover, the lakes support recreational fishing and other tourism-related activities.

The four lakes that we studied were selected based on their sources of walleye recruitment (natural or stocked) and their environmental status (degraded or non-degraded). The location of the lakes in the Clay Belt (Figure 1a) offered a unique opportunity to evaluate the response of stocked walleye populations in their preferred yet degraded environments. Lake Osisko (N48.24, W79.00) (referred as D1) and Lake Dufault (N48.31, W79.00) (referred as D2) are both degraded and were historically stocked with walleye. D1 was stocked between 1999 and 2018 (Table S1), and D2 between 1986 and 2015 (Table S2). Stocking efforts have therefore been completed for four years in D1 and seven years in D2 prior to this study. Consequently, adult walleye captured during this study could have been either stocked or naturally recruited, while young-of-the-year and juvenile captures would have been naturally recruited.

**Figure 1.**
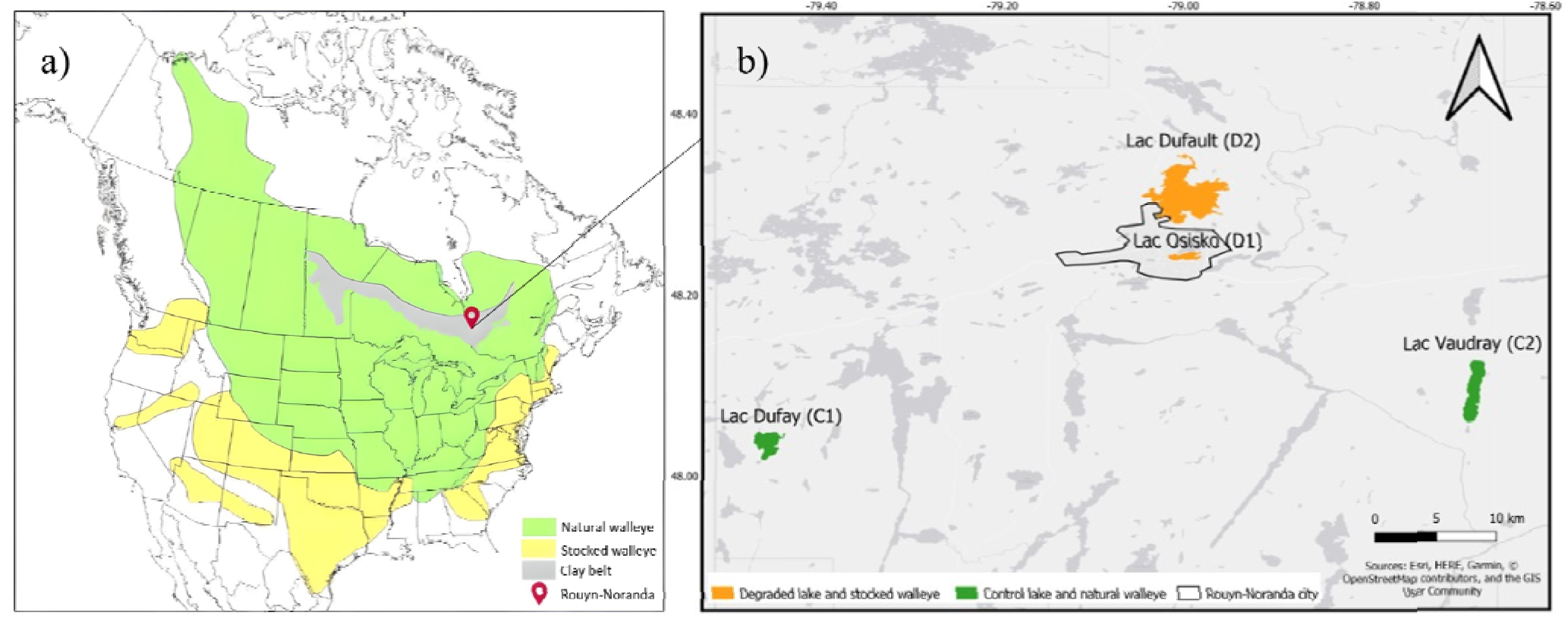
a) Natural and introduced distribution range of walleye in North America. The Clay Belt is indicated in grey (adapted from Hartman (2009)). b) Study sites in the Abitibi-Témiscamingue region near Rouyn-Noranda (Quebec, Canada).

The stocked and degraded lakes were compared to Lake Dufay (N48.04, W79.46) (referred as C1) and Lake Vaudray (N48.10, W78.68) (referred as C2) (Figure 1b), both of which support natural walleye populations. C1 is relatively pristine, exhibiting minimal metal contamination and few anthropogenic disturbances. However, C2, which lies in the path of prevailing winds coming from Rouyn-Noranda, has an intermediate level of metal in the regional contamination gradient (Laflamme et al. 2000). C2 is located in the Joannès-Vaudray biodiversity reserve and has permanent residences and summer cottages on its eastern shore (Gouvernment du Quebec 2015).

### Habitat characterization

In each lake, three pelagic stations were sampled, including the deepest point of the given lake. Habitat requirements for larvae, young-of-the-year, juveniles, and adults were characterized by sampling on five occasions, i.e., in spring (June 2022), in summer (August 2022), in fall (September 2022 and 2023), and in winter (March 2023).

At each station and each sampling occasion, water pH, dissolved oxygen concentration (mg/L) and saturation (%), temperature (°C), and specific conductivity (µS/cm) were measured with a multi-parameter probe through the water column (RBR Concerto, Ottawa, Canada). Light penetration was assessed with a light sensor (LI-1500, Li-Cor, Lincoln, USA) to determine the euphotic zone. Light measurements were taken at water surface, 0.5 m and then at every meter of the water column until 1% of incident surface light was measured. Lake water was collected every meter of the epilimnion with a Van Dorn or Ruttner bottle and mixed in a cleaned bucket. From this lake water, two replicates of 40 mL water samples were collected into 50 mL acid-washed vials (10% HCl) for total nitrogen and total phosphorus analyses.

Based on data collection during seasonal water column sampling (winter, spring, summer, and fall) of the water column, the annual mean, maximum, and minimum values of pH, dissolved oxygen concentration (mg/L), specific conductivity (µS/cm), temperature (°C), and concentrations of phosphorus (mg/L) and nitrogen (ppm) were calculated. Dissolved organic and inorganic matter sampled during the fall of 2022 was also included in the descriptive analysis.

All data and statistical analyses were conducted using R software version 4.3.3 (R Core Team 2024). To compare photic zones among lakes and seasons (spring, summer, and fall), considering their interaction, an analysis of variance (ANOVA) and Tukey test were performed. ANOVA’s assumptions were tested using residual plots and formal tests (Shapiro-Wilk test for normality and Levene’s test for homogeneity of variance).

### Characterization of walleye populations

Young-of-the-year were sampled at night during September 2023 using a five-meter aluminum boat equipped with a generator powered pulsator (GPP model 2.5 from Smith-Root) (Paquin and Brodeur 2021). Transects were conducted for 20 min along shorelines until 22 to 30 young-of-the-year individuals were collected. The smallest length-class encountered in each lake (Table S3) were captured to maximize the likelihood to collect young-of-the-year individuals, and their age was later confirmed by sagittal otolith reading. The transects were selected following these criteria: < 2 m deep, gently sloping banks, and proximity to aquatic vegetation. Bathymetric maps (Navionics) and satellite images (Google Earth) were consulted for site selection.

Juvenile and adult walleye were collected during standardized fishing surveys conducted by the *Ministère de l’Environnement, de la Lutte contre les changements climatiques, de la Faune et des Parcs* (MELCCFP) in September 2021 (C1), 2022 (D1, C1, and C2), and 2023 (D2) (Service de la faune aquatique 2011). Eight-panel fishing gillnets that were 7.6 m long and 1.8 m in height were deployed in order of increasing mesh size (25, 38, 51, 64, 76, 102, 127, and 152 mm). Nets were placed horizontally at depths of 2–15 m and were left in place for 18–24 h, always including the night. The number of nets used was 8 in D1, 10 in D2 and C1 lakes and 11 in C2.

For each walleye encountered in this study, total length and body mass were measured, and sex and sexual maturity were compiled. Sexual maturity was determined based on observation of gonadal development: individuals exhibiting advanced development are expected to participate in the next spawning event (Duffy et al. 2000). Muscle samples were extracted from behind the dorsal fin and frozen at −80 °C. Sagittal otoliths were also collected from all walleye individuals. All fish body measurements and extractions were completed within 24 h after the sampling. Right sagittal otoliths were embedded in epoxy (Miapoxy 100 resin and 95 hardener) and sliced vertically using a low-speed saw (Buehler^®^ IsoMet^®^) for age reading of individuals. Otoliths were aged twice by independent observers.

We followed the UQAT animal ethics protocol during the handling and euthanasia of net-caught fish and obtained a permit (2022-07-19-066-08-SP) from the MELCCFP.

#### Analysis

##### Walleye relative abundance

Walleye were categorized by life stages: young-of-the-year, juvenile and adult. The boundary between juveniles and adults was set at the age at which 50% (A50) of the individuals in a population are expected to exhibit developed gonads and thus, participate in the next spawning event (Mainguy et al. 2024). Using the method detailed in Mainguy et al. (2024), the A50 was estimated using generalized linear models (GLMs) with three different link functions (logit, probit, cloglog). Model fit was assessed using the McCullagh and Nelder (1989) and Osius and Rojek (1992) goodness-of-fit tests, and models with adequate fit were then compared using Akaike Information Criterion (AIC, cutoff of 2).

The number of walleyes captured in each age category was standardized by fishing effort. Young-of-the-year involved 20-min electrofishing transects (number of young-of-the-year/20-min transect). The number of juveniles and adults caught during 18–24 h of gillnet fishing were standardized to the expected catch for a hypothetical 24-h net-set period. Young-of-the-year captured in gill nets were excluded from the analysis.

Catch-per-unit-effort (CPUE) for each life stage was compared between lakes using GLMs (negative binomial distribution) using the *MASS* package (Venables and Ripley 2002). The goodness-of-fit was tested with the *hnp* package (Moral et al. 2017; MainGuy and Moral 2021). A post-hoc test was realised on the best model with the *emmeans* package to compare the relative abundance between lakes (Lenth R 2024).

##### Walleye growth

The growth of walleye in each lake was modeled using the Von Bertalanffy growth function (VBGF) from the *FSA* package (Ogle et al. 2023; Equation 1):

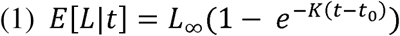

Where *E*[*L*|*t*] = expected or average length at age t;*L*_∞_= Asymptotic average length of sampled population; K = Brody growth rate coefficient (year^-1^); t_0_ = modeling artifact that is said to represent the time or age when the average length was zero (Ogle et al. 2023). Parameters were pre-optimized using the *vbStart* function (Ogle et al. 2023). Confidences intervals for each VBGF model were obtained using the *investr* function to compare mean length-at-age in each lake (Greenwell and Kabban 2014).

### Evaluation of lower trophic levels biomass and diversity

Food resources for larvae were evaluated by sampling zooplankton communities once in each lake. Sampling occurred between 30 May and June 2, 2023. The timing of sampling was determined according to water surface temperatures following ice-outs. Ice-out dates and water temperatures were assessed once a day by volunteers in accessible lakes: D1, D2 and C2. In remote C1, ice-out date was photo-interpreted with NASA Worldview (NASA 2025) and temperature continuously measured using a temperature data logger (iButton, Alphamac Inc., DS1922L, Ste-Julie, Canada) near the shore. The sampling of zooplankton was conducted 3 – 8 days after the expected end of egg hatching i.e. 210 temperature unit with water at 8 – 15°C (Bruce et al. 2011), which allow larvae to consume yolk sac and start to feed on zooplankton (Engel et al. 2000). Zooplankton sampling therefore occurred 29 – 31 days after ice-out.

Vertical tows were conducted through the water column using a 50 µm mesh net of 25 cm diameter. Three stations per lake were selected based on known (Ressources Falco, 2018; M. Bélanger, MELCCFP, Rouyn-Noranda, Canada, personal communication, 2023) or potential spawning sites (Navionics and satellite image). Sampling was conducted in pelagic areas (2 – 12 m), perpendicular to spawning sites, because walleye larvae are thought to have poor swimming abilities and to feed in pelagic areas near spawning sites once yolk sac is absorbed (Faber 1967, Hartman 2009). Stations were selected to cover spatial heterogeneity and zooplankton communities representative of the pelagic zone of the lakes (Beisner et al. 2006, Grosbois et al. 2017). Two 20 L water column replicates were collected at each site (n = 6 per lake) using a Van Dorn or Ruttner bottle, sieved through a 50 µm net, and preserved in 95% ethanol. Macroinvertebrate community samples were collected during July 2022 for identification and biomass measurement. Three replicates in each of the three littoral sites (n = 9 per lake) were collected using a kick-net. Samples were collected from representative lake substrates (mainly sand, sediments, and pebbles), accounting for habitat diversity including proximity to aquatic plants, and at depth < 1 m in the littoral zone using the swirling method for 30 s and a collection area of ∼0.2 m^2^. Macroinvertebrates samples were immediately preserved in ethanol 95%.

Food resources for juvenile and adult walleye were evaluated by sampling forage fish communities during July 2022 using gillnets (10 m length, 1 m height and 10 mm mesh size). Gillnets were positioned perpendicular to the shore at depths of 0.5–5 m (n = 3 per lake) and left in place for 18 – 24 h. For each fish captured, specie identification, total length and body mass were measured. Muscle samples were dissected from behind the dorsal fin were frozen at −80 °C.

Zooplankton (from 20 L water column samples, May 2023) and macroinvertebrates samples (from kick-net samples, July 2022) preserved in ethanol 95% were counted and sorted to the family level, except the female and copepodite from the Calanoida order that were identified to order level. The entire sample was counted and identified, or split when abundances were too high (Blackburn-Desbiens et al. 2023). The total lengths of the first 30 individuals of each family were measured using Zeiss Zen 3.9. software and a camera (AxioCam 208 color, Zeiss, Oberkochen, Germany) mounted on a stereomicroscope (Zeiss discovery V12, Oberkochen, Germany).

Total length was used to estimate biomass of zooplankton and macroinvertebrates using published length-mass regressions (Smock 1980, Rowe and Berrill 1989, Benke et al. 1999, EPA Great Lakes National Program Office 2003, Méthot et al. 2012). Fish biomass was estimated by accounting for the average water loss per fish (82%) during the freeze-drying process, a loss that was consistent across species. ANOVAs with Tukey’s test were conducted to compare the biomass of zooplankton (µg/L), macroinvertebrates (mg/kick-net), and fish (g/10 mm gill net standardized to 24 h) among lakes. The biomass of macroinvertebrate orders was log transformed to meet the assumption of normality of residuals for ANOVA. Bycatches (fish with total length > 30 cm) were excluded from the analysis.

Taxonomic diversities were estimated with Shannon’s diversity index (H’) on adult and copepodites when identified to family level, excluding nauplii (Wiener et al. 1950; Hasan et al. 2023). Shannon’s index was calculated based on the proportion (*n_i_/N*) of each taxon found in each sampling unit (pelagic station, kick-net, or 10 mm gill net) relative to the total number of organisms counted (*N*) (Equation 1):

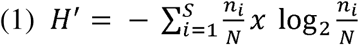

### Food web Sstable isotopes and stomach contents

Food resources for larval walleye were assessed with vertical tows conducted through the water column using a 50 µm mesh net of 25 cm diameter in May 2023.

To assess food resources for young-of-the-year, macroinvertebrate communities in the littoral zone were sampled one week before electrofishing using a 350 µm mesh kick-net. Three kick-net replicates from the orders Amphipoda, Coleoptera, Diptera, Ephemeroptera, Odonata, and Trichoptera were collected for stable isotope analysis. We also collected ten individuals of the most abundant fish species (other than walleye) collected during electrofishing. Fish smaller than young-of-the-year walleye were selected to represent potential prey (Table S4).

Zooplankton, littoral macroinvertebrates and fish were sorted and frozen to −80 °C within 24 h of the sampling. Zooplankton samples were kept in the dark at 4 °C and were passed through stacked sieves (500, 355, 250, 106, or 63 μm mesh), placed from coarse to fine.

Zooplankton, macroinvertebrates, and fish samples were freeze-dried (Pfeiffer Vacuum, D-35614, Asslar, Germany), ground to a fine powder, and encapsulated in 5 x 3.5 mm tin capsules for stable isotope analyses (Grosbois et al. 2017, 2020). Samples of 1.0 ± 0.1 mg dry weight were measured on a microbalance (Geneq Inc. Model # AP250D), and their N and C isotopic composition was analyzed using an Elementar Pyrocube elemental analyzer interfaced to a Delta V Plus IRMS mass spectrometer. Replicate analyses of isotopic standard reference materials USGS 40 (δ^13^C = –26.39‰; δ^15^ N = –4.52‰) and USGS 41 (δ^13^C = 37.63‰; δ^15^ N = 47.57‰) were used to normalize isotopic values of working standards to the Air (δ^15^ N) and Vienna Pee Dee Belemnite (δ^13^C) scales.

Isotope values are expressed in δ notation following the formula δX (‰) = [(Rsample/Rstandard) – 1] × 103, where X is ^13^C or ^15^N and R is ^13^C/^12^C or ^15^ N/^14^N isotopic ratio. Working standards were analyzed after every ten samples to monitor instrument performance and check data normalization. The precision of the laboratory standards was ±0.3‰ for C and N.

The isotopic data of δ^15^N and δ^13^C were categorized by taxonomic group. Three replicates were analyzed for each zooplankton sieve and macroinvertebrates orders, sixty for walleye (30 randomly selected mixed-age fish and 30 young-of-the-year), and 10 for other fish species. Walleyes were divided into length classes: 8–13 cm, 14–30 cm, and > 30 cm.

We used a two-way ANOVA with Tukey’s test to compare walleye δ^15^N and δ^13^C muscle composition by length classes among lakes. We also used permutational ANOVAs (PERMANOVAs) to compare the δ^15^N and δ^13^C values for each taxonomic group. This analysis was done to describe the food web of each lake. The PERMANOVAs were performed with 999 permutations on the distance matrix (Euclidean distances; δ^15^N and δ^13^C combined or singularly) with the R packages *vega*n (Oksanen J. et al. 2024). Post-hoc pairwise tests were performed using *pairwiseAdonis* package (Martinez Arbizu 2020).

#### Stomach contents

Sixty stomachs (30 young-of-the-year and 30 juveniles or adults) were extracted from fish individuals within 24 h of sampling and preserved in 95% ethanol in order to minimize the digestion of prey items, especially invertebrates and soft bodied fishes that can be digested rapidly (Bowen 1996). They were then dissected using a binocular microscope and prey items were counted and identified to species (fish) or family (invertebrates) levels.

Walleye stomachs were separated by length class for diet analysis. For each lake and length class, prey items were counted, identified, and expressed as percentages. Empty stomachs were excluded from the analysis. Categories such as “Fish remains” and “Invertebrate remains” refer to stomachs where fish or invertebrates were the only prey items but could not be further identified.

## Results

### Habitat characterization

In regard to the water quality, the annual mean (i.e., all four seasons combined) of specific conductivity was higher in degraded lakes (ANOVA, F_(3,14)_ = 2885, p < 0.0001 (Table 1), particularly in D1 (Tukey, p < 0.0001). On average, the specific conductivity was 10-times higher in D1 and 4-times higher in D2 compared to C1 and C2 lakes. In D1, pH values exceeding 10 were recorded in summer and were higher than the other three lakes throughout all seasons (ANOVA, F_(3,14)_ = 34.9, p < 0.001). The mean depth of the photic zone (< 1% ambient light) was greater in degraded lakes (ANOVA, F_(3,42)_ = 117.4, p < 0.0001). Photic zones were significantly larger in degraded lakes in comparison to the non-degraded lakes (Tukey, p < 0.05), for each season except of fall, where the photic zone in C2 was slightly deeper than during the other seasons. The mean depth of photic zone ranged from 3.8 m (± 0.3; D2, fall) to 5.4 m (± 0.9; D1, fall) in degraded lakes and from 2.0 (± 1.0; C1, spring) to 3.0 m (± 0.0; C2, fall) in the two control lakes.

**Table 1.** Mean, minimum, and maximum of measured habitat variables for all four seasons, except DOC and DIC, which were only measured in the fall in the four study lakes. Variables were taken in the water column (specific conductivity, dissolved oxygen, temperature, and pH) or epilimnion (DOC, DIC, phosphorus, and nitrogen concentrations).

| Habitat variables | -----Study Lakes----- |  |  |  |
| --- | --- | --- | --- | --- |
|  | Osisko (D1) | Dufault (D2) | Dufay (C1) | Vaudray (C2) |
| Location | 48.2412,<br>-79.01308 | 48.30709,<br>-78.98737 | 48.04098,<br>-79.45187 | 48.09551,<br>-78.67839 |
| Lake Area<br>(km<sup>2</sup>) | 4.76 | 28.48 | 5.27 | 8.63 |
| Maximum depth<br>(m) | 8 | 19.5 | 12 | 32 |
| Secchi depth<br>(m) | 2.80<br>[2.35–3.35] | 2.58<br>[2.15–3.23] | 1.07<br>[0.80–1.41] | 1.60<br>[1.15–1.98] |
| Photic zone ($< 1\%$<br>ambient light; m) | 5.1<br>[4.0–6.0] | 4.2<br>[3.5–5.0] | 2.1<br>[2.0–3.0] | 2.4<br>[1.0–3.0] |
| Specific<br>conductivity<br>($\mu\text{S}/\text{cm}$) | 284.2<br>[277.0–362.5] | 125.8<br>[108.2–267.9] | 28.9<br>[23.3–95.9] | 27.4<br>[23.1–132.8] |
| Dissolved oxygen<br>(mg/L) | 9.4<br>[1.4–13.3] | 8.2<br>[0.04–12.6] | 7.3<br>[0.6–13.3] | 8.5<br>[0.2–14.0] |
| Temperature<br>(°C) | 17<br>[0.3–22.2] | 15.6<br>[0.2–21.6] | 17<br>[0.2–22.1] | 12.8<br>[0.2–22.2] |
| pH | 8.63 | 7.70 | 7.39 | 7.04 |
|  | [6.95–10.68] | [6.43–8.40] | [5.78–8.43] | [5.68–8.30] |
| Total phosphorus | 17.9 | 12.4 | 22.3 | 8.1 |
| (µg/L) | [14.2–21.0] | [9.9–16.7] | [21.4–23.1] | [7.6–8.5] |
| Total nitrogen | 0.285 | 0.244 | 0.226 | 0.215 |
| (ppm) | [0.226–0.356] | [0.180–0.395] | [0.188–0.298] | [0.175–0.255] |
| DIC | 11.59 | 5.71 | 2.31 | 1.92 |
| (mg/L) | [10.84–12.11] | [5.35–6.31] | [2.05–2.57] | [1.57–2.42] |
| DOC | 2.38 | 6.28 | 10.29 | 10.45 |
| (mg/L) | [2.76–2.87] | [6.04–6.50] | [9.92–10.67] | [10.04–11.04] |

### Walleye relative abundance and A50

Walleye matured younger in degraded lakes. For males, A50 was 3 and 4 years in D1 and D2 lakes, while it was 6 years in control lakes. Females matured at older ages than males in all lakes: A50 was 5 and 6 years old in D1 and D2, while it was 9 years in control lakes (Table S5).

Young-of-the-year CPUE were higher in degraded lakes than in control lakes: the highest CPUE for young-of-the-year was recorded in D1 (mean ± SE: 27.6 ± 13.0 fish/20 min; GLM, Z_(3,37)_ = 14.5; p < 0.0001), followed by D2 (7.4 ± 3.6 fish/20 min; GLM, Z_(3,37)_ = 10.4; p < 0.0001) (Figure 2a). The young-of-the-year relative abundance between control lakes were similar, with 2.0 ± 1.7 fish/20 min in C1 and 1.8 ± 1.7 fish/20 min in C2 (*emmeans* contrast, Z.ratio = 0.398; p = 1.0).

**Figure 2.**
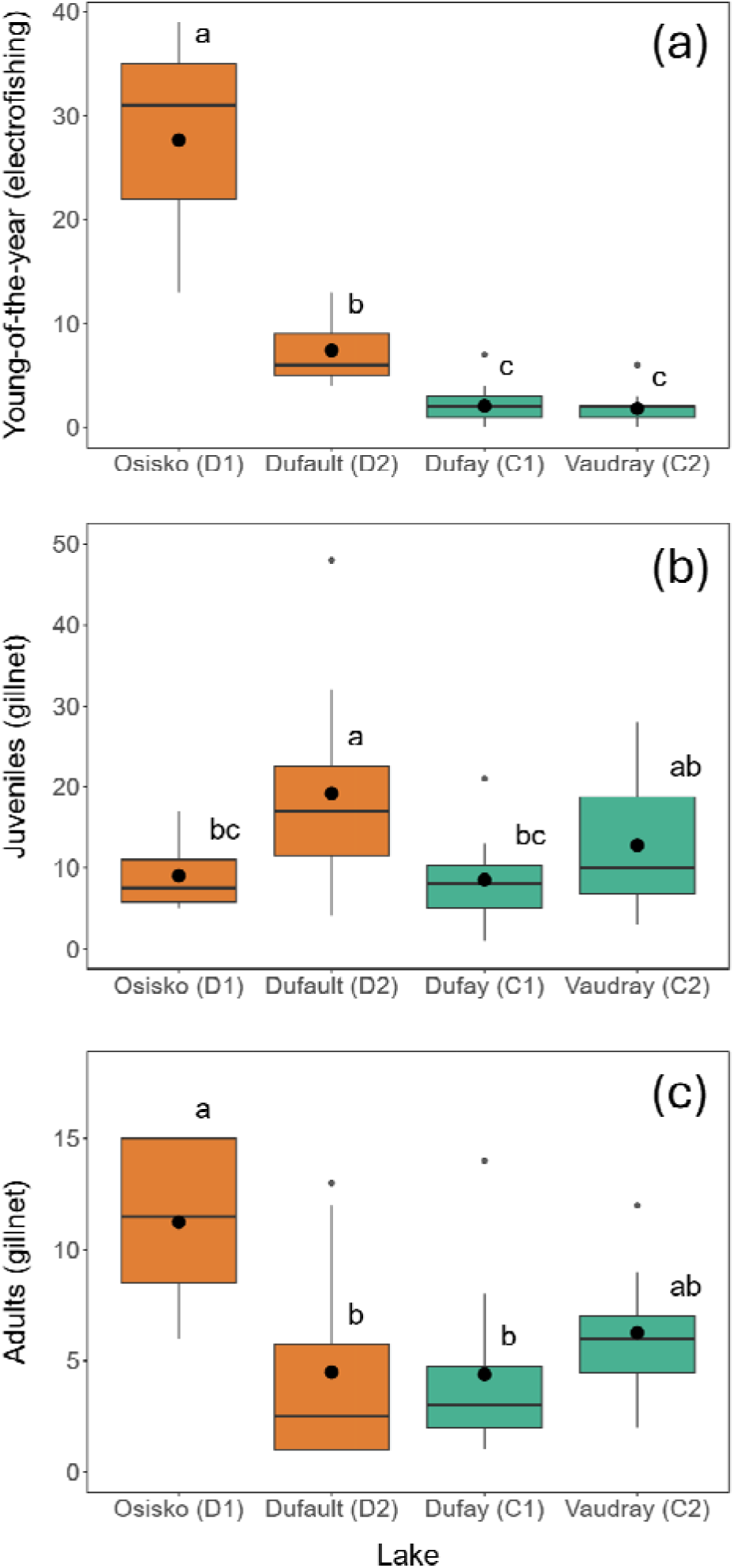
Catch-per-unit-effort (CPUE) in the four study lakes. Black dots indicate means, and letters represent significant differences among lakes (Tukey tests). (a) Young-of-the-year standardized to captures per 20 min electrofishing transects, (b) Juveniles (individuals captured/24 h gillnet sets), aged based on A50 results for lake and sex, (c) Adults (individuals captured / 24 h gillnet sets), aged based on A50 results for lake and sex.

Catch-per-unit-effort of juvenile captured per 24 h using gill nets was greater in D2 (19.2 ± 12.3 fish/24h gillnet) than in D1 (9.0 ± 4.4 fish/24h gillnet; *emmeans* contrast, Z.ratio = −2.8; p = 0.03) and C1 (8.5 ± 5.1 fish/24h gillnet; *emmeans* contrast, Z.ratio = 0.2; p < 0.01), but not C2 (12.8 ± 8.3 fish/24h gillnet; *emmeans* contrast, Z.ratio = 1.8; p = 0.3) (Figure 2b). No other pairwise differences were recorded (p > 0.05).

The number of adult walleye captured per 24 h of gill net fishing was greater in D1 (11.3 ± 3.9 fish/24h gillnet) than in D2 (4.5 ± 4.6 fish/24h gillnet; *emmeans* contrast, Z.ratio = 3.3; p < 0.01) and C1 (4.4 ± 3.9 fish/ 24h gillnet; *emmeans* contrast, Z.ratio = 3.3; p < 0.01) (Figure 2c). Adult relative abundance was similar between D2, C1 and C2 (6.3 ± 2.7 fish/24 h gillnet; *emmeans* contrast, Z.ratio = −1.3–0.1, p > 0.5).

The mean age of adults was lower in degraded D1 (5.4 ± 0.8) and D2 (5.3 ± 1.6) lakes than in control C1 (8.5 ± 1.8) and C2 (9.0 ± 1.5) lakes (Tukey, p < 0.001).

### Walleye growth

Growth was rapid in degraded lakes, with body growth coefficient (K) of 0.24 year^-1^ (D1) and 0.29 year^-1^ (D2), which are approximately 2.4 to 3.9 times higher than those observed in control lakes: 0.07 year^-1^ (C1) and 0.10 year^-1^ (C2) (Figure 3). Autumnal young-of-the-year were the longest in D1, followed by young-of-the-year in D2, in C1, and finally in C2 (ANOVA, F_(3, 116)_ = 125.9, p < 0.001). At age 0+, walleye from D1 measured 14.5 cm ± 1.1 (mean ± SE), which was comparable to the size of walleye from C2 at age 1 (14.7 cm ± 1.8, n = 27).

**Figure 3.**
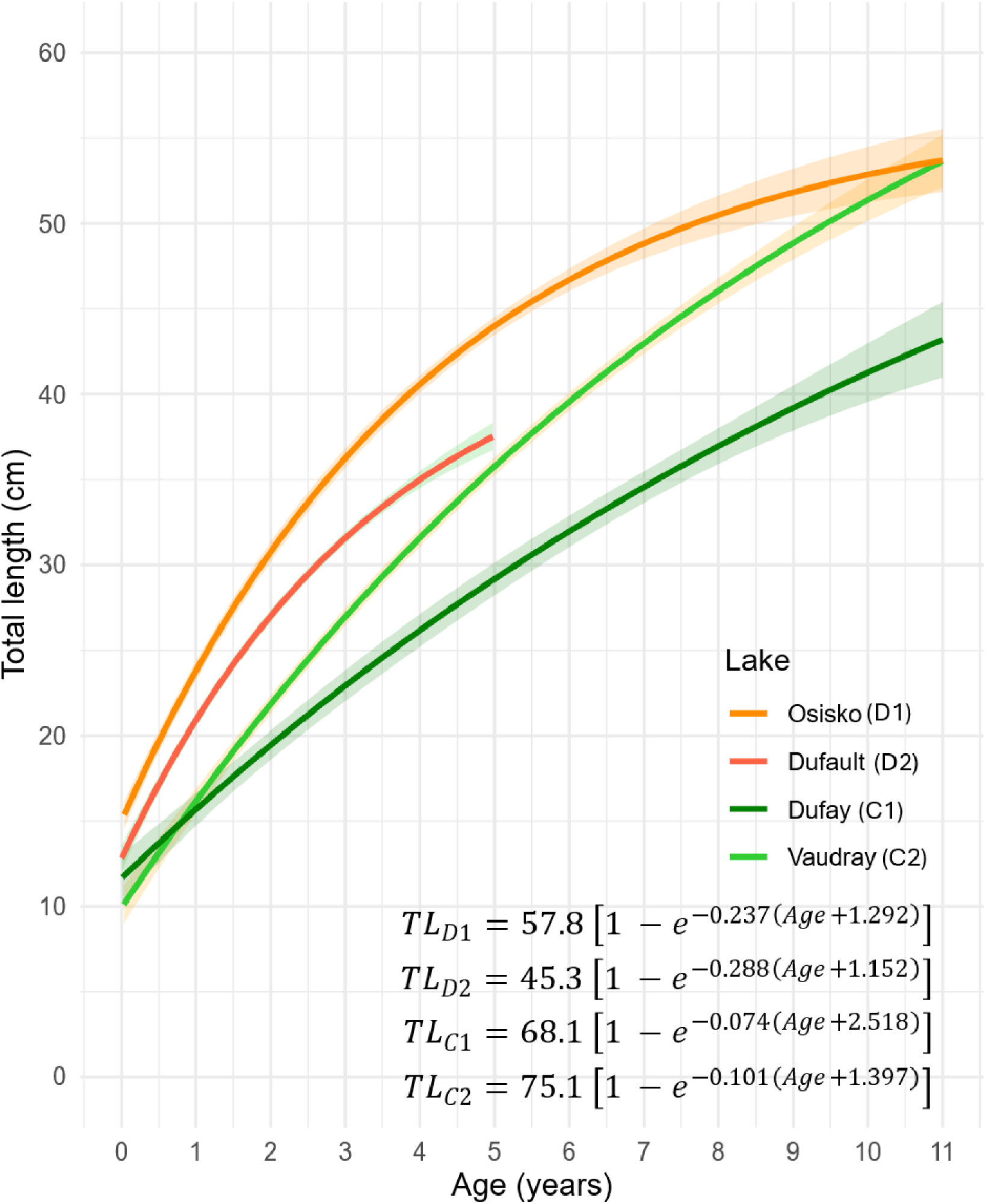
Von Bertalanffy walleye growth regression models, 95% confidence interval for D1 (n = 186), D2 (n = 288), C1 (n = 167), and C2 (n = 270), together with their VBGF equations.

Despite the fast growth at a young age, the asymptotic average length (L_inf_) was lower in degraded lakes—57.8 cm (D1) and 45.3 cm (D2)—than in control lakes: 68.1 cm (C1) and 75.1 cm (C2). Furthermore, only 4 % (D1, n = 158) and 1 % (D2, n = 264) of walleye caught in gillnets were 10 years old or older in degraded lakes compared to 13 % (C1, n = 137) and 15 % (C2, n = 242) in control lakes. The maximum sampled age was also lower in degraded lakes: 11 (D1) and 14 years old (D2), compared to control lakes: 18 (C1) and 20 years old (C2).

### Food resources

#### Zooplankton

Spring zooplankton biomass was higher in D2 (166.1 ± 40.5 μg/L) than in C1 (74.4 ± 49.3 μg/L; Tukey, p = 0.03) and C2 (29.2 ± 7.9 μg/L; Tukey, p < 0.001) (Figure 4a). Zooplankton biomass in D1 (102.0 ± 81.6 μg/L) during spring did not differ from the other three lakes (Tukey, p > 0.1). Zooplankton families were more diversified in D2 (1.5 ± 0.17; Tukey, p < 0.001) and C1 (1.6 ± 0.3; Tukey, p < 0.005) than in D1 (0.4 ± 0.2) and C2 (0.9 ± 0.2; Figure 4b). The Shannon diversity index (H’) was greater by 0.5 (Tukey, 95% CI: 0.1–0.8) in C2 than in D1 (Tukey, p < 0.01).

**Figure 4.**
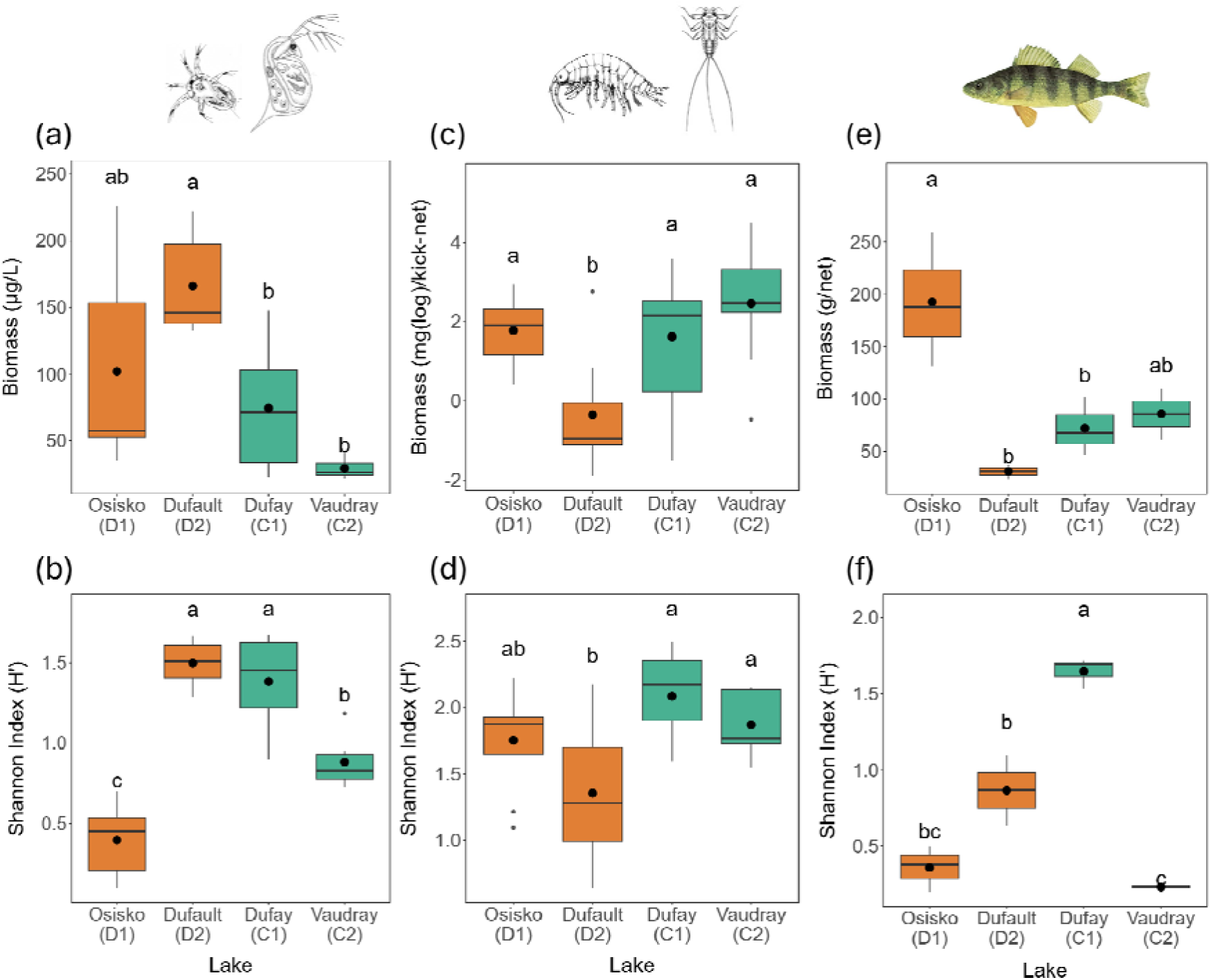
(a and b) Biomass and Shannon diversity index of zooplankton in spring, (c and d) macroinvertebrates in July 2022, and (e and f) forage fish in July 2022 in the four study lakes.

The characteristics of the zooplankton communities differed between degraded and control lakes. Copepods were the most common order in all lakes, but Cyclopidae dominated degraded lakes while Calanoida dominated control lakes (Figure S1). Daphniidae were also more abundant in degraded lakes (25.7 ± 24.7 µg/L D1; 41.0 ± 12.7 µg/L D2) than in control lakes (5.7 ± 4.7 µg/L in C1; 0.50 ± 0.0 µg/L in C2) (Figure S2).

#### Macroinvertebrates

Macroinvertebrate biomass per kick-net sample was significantly lower in D2 than in the other three lakes (ANOVA, F_(3, 36)_ = 6.6, p = 0.001) (Figure 4c). The abundance of macroinvertebrates was similar in the other lakes (Tukey, p < 0.5).

Shannon’s H’ for macroinvertebrate orders was lower in D2 (1.3 ± 0.5) than C1 (2.1 ± 0.3; p = 0.001) and C2 (1.9 ± 0.2; p = 0.03) (Figure 4d). Although the biomass and diversity of macroinvertebrates were similar in D1 (1.8 ± 0.4) (Tukey, p > 0.1), C1 and C2, their community composition differed. Ephemeroptera, Plecoptera, and Megaloptera were absent from samples in degraded lakes (D1 and D2). Pooling all samples (n = 9 in each lake), the taxonomic richness was lower in degraded D1 (13) and D2 (11) lakes than in control C1 (21) and C2 (22) lakes.

#### Fish

The total dry biomass of forage fish was significantly higher in D1 (192.8 ± 63.7 g/net) than in D2 (31.1 ± 10.0 g/24 h; Tukey, p = 0.02) and C1 (72.2 ± 28.1 g/net; Tukey, p = 0.05) (Figure 4e). Only six forage fish individuals were captured in D2. Total biomass in C2 (86.1 ± 24.4 g/net) was similar to that in the other three lakes (Tukey, p > 0.1).

With ten forage species recorded, including the shiners, golden shiner (*Notemigonus crysoleucas*), common shiner (*Luxilus cornutus*), spottail shiner (*Notropis hudsonius*), and mimic shiner (*Notropis volucellus*), C1 (1.6 ± 0.1) had higher Shannon index than D1 (0.4 ± 0.2), D2 (0.9 ± 0.3), and C2 (0.2 ± 0.0) lakes (Tukey, p < 0.01) (Figure 4f). In D1, walleye and yellow perch (*Perca flavescens*) were captured, and yellow perch biomass in this lake was greater than in C1 (Tukey, p < 0.02) and Dufault (p < 0.01)).

### Food web stable isotopes and stomach contents

Walleye trophic level was highest in D1 (δ^15^N = 13.6 ± 0.9), followed by D2, (δ^15^N = 10.1 ± 1.0), C1 (δ^15^N = 9.7 ± 0.8), and C2 (δ^15^N = 8.1 ± 0.9; Tukey, p < 0.0001). Total length was positively correlated with δ^15^N for all lakes (Pearson correlation = 0.397, p < 0.0001). Moreover, significant interactions between fish length classes and lake were recorded for δ^15^N (ANOVA, F_(6, 215)_ = 3.3, p = 0.004) and δ^13^C (ANOVA, F_(6, 215)_ = 3.2, p = 0.005).

Carbon isotope ratios were higher in D1 walleye in length classes 8–13 cm (δ^13^C = −24.1 ± 2.5) and 14–30 cm (δ^13^C = −24.5 ± 2.7) than in fish from the other three lakes (Tukey, p < 0.05). Walleye > 30 cm from D1 (δ^13^C = −25.0 ± 1.6) also had higher δ^13^C values, except when compared with smaller walleye from the same lake and 8–13 cm walleye from C1 (δ^13^C = −27.1 ± 1.0) (95 CI: −0.02–4.4; Tukey, p = 0.06). Carbon isotope rations in were similar in the remaining lakes and fish length classes (Tukey, p > 0.05).

#### Trophic relationships within lakes

Trophic relationships between different taxonomic groups of organisms, including walleye, differed in each lake (Figure 5).

**Figure 5.**
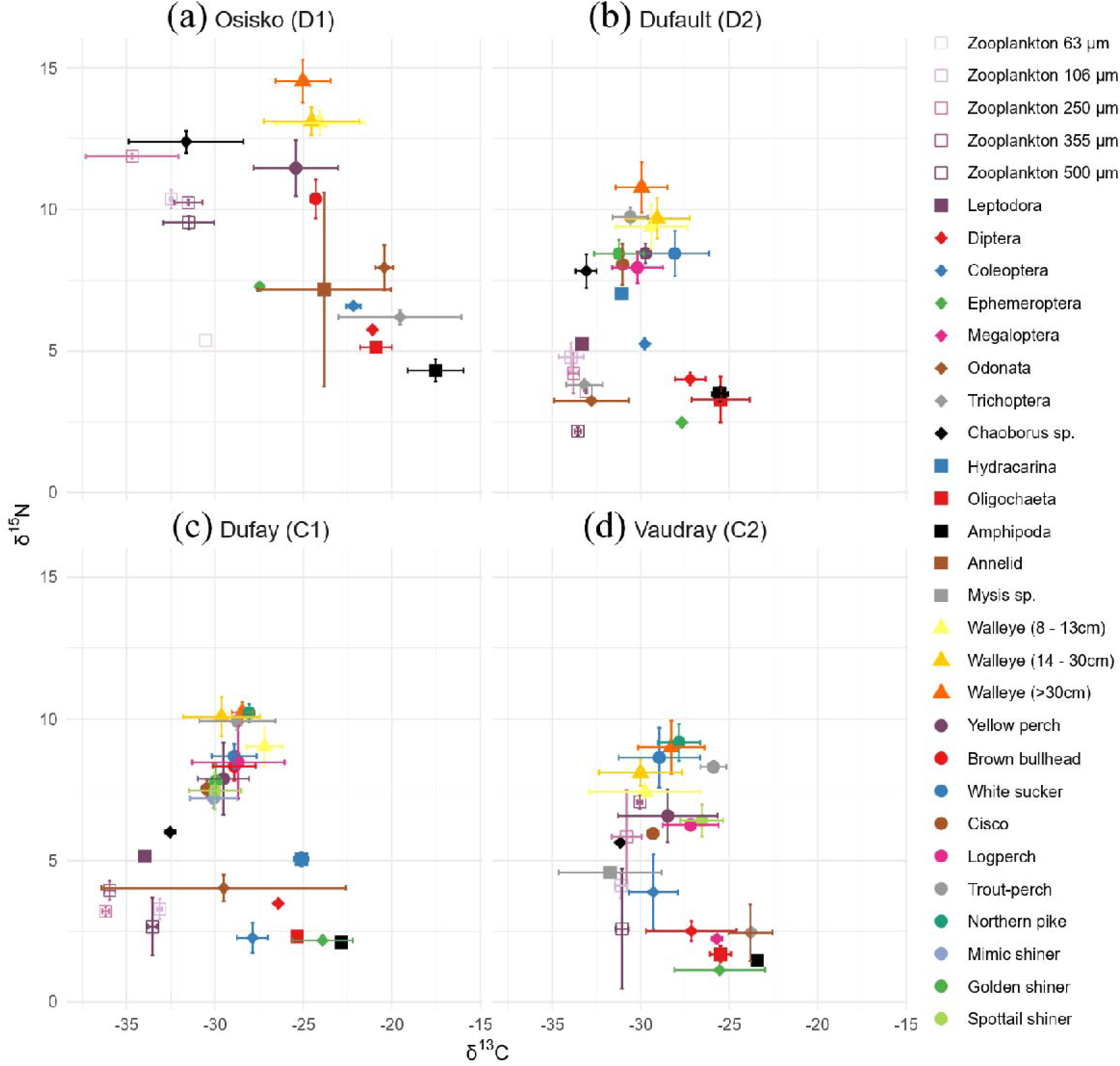
Stable isotope values δ^15^N and δ^13^C in (a) Osisko (D1), (b) Dufault (D2), (c) Dufay (C1), and (d) Vaudray (C2) lakes (means ± standard deviation). Zooplankton dominate the bottom left of each chart (low trophic level and pelagic), while macroinvertebrates dominate the bottom right (low trophic level and benthic). Fish occupied higher trophic levels and were distributed across the δ^13^C axis according to their diet.

#### D1 (Osisko Lake)

All size classes of walleye had higher δ^15^N values than macroinvertebrates groups, zooplankton and other fish species (pairwise, F = 4.3–345.5, p < 0.05, Figure 5a). Walleyes > 30 cm long had higher δ^15^N values than smaller size classes (pairwise, F = 25.8–57.8, p < 0.01) but similar δ^13^C values (pairwise, F = 1.8–2.7, p > 0.05). Walleye 8– 13 cm and 14–30 cm had similar δ^15^N and δ^13^C (pairwise, F = 0.06, p = 0.98). The main prey items consumed by these size classes were yellow perch (4.0–22.2 cm) and annelids (pairwise, F (δ^15^N) = 14.0–58.7, p < 0.01; F (δ^13^C) = 1.0–2.2, p > 0.11). Yellow perch or fish remains constituted 71% (8–13 cm, n = 6), 83% (14–30 cm) (n = 35) and 62% (> 30 cm, n = 15) of walleye diets (Figure 6). Plants were also found in stomachs, especially frequent in those of the > 30 cm size category (19%). Plants were absent in walleye stomachs sampled from the other three lakes.

**Figure 6.**
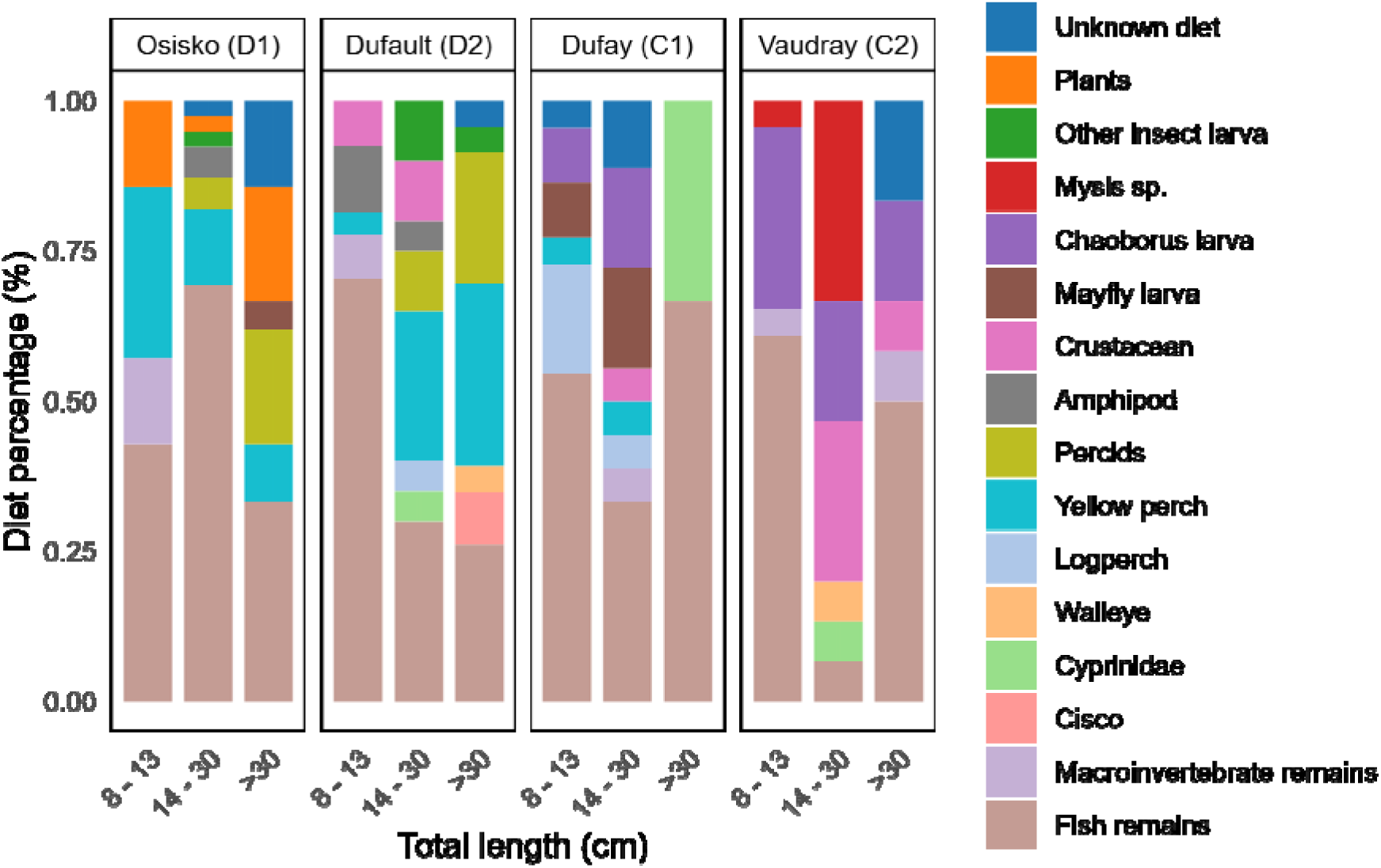
Stomach contents of walleye in 8–13 cm, 14–30 cm, and > 30 cm size classes. Diet percentages (%) are based on the frequency of prey items relative to the total number of prey items found per lake and size category. Empty stomachs were excluded from analysis.

#### D2 (Dufault Lake)

Walleyes > 30 cm were top predators and had higher δ^15^N values than all other organisms (pairwise, F = 12.3–121.8, p < 0.01, Figure 5b). Smaller walleye size classes had similar δ^15^N and δ^13^C (pairwise, F = 0.4, p = 0.8). Smaller size classes were not at the top of the food web, and their δ^15^N values were similar to those of trout-perch (*Percopsis omiscomaycus*; 5.2–10.2cm) (pairwise, F = 0.9–2.2, p > 0.05). Yellow perch was the most frequent fish species in walleye stomach from all size classes (4–30%, n = 54). Walleye > 30 cm had similar δ^13^C values than smaller individuals (pairwise, F = 1.0–2.4, p > 0.1). Cisco (*Coregonus artedi*) constituted 9% of > 30 cm walleye diet (n = 17, Figure 6). As in D1, fish constituted the major prey items in every walleye length class (77–89%).

#### C1 (Dufay Lake)

In contrast to their top predator status in D1 and D2 (Figure 5c), the trophic status of walleye in C1 was size dependant and modulated by the presence of northern pike (*Esox lucius*). Northern pike (38.6–62.5 cm) had higher δ^15^N values than walleye 8–13 cm (pairwise, F = 8.4, p < 0.01) and similar δ^15^N to walleye 14–30 cm and > 30 cm (pairwise, F = 0.5, p > 0.5), suggesting that this species competed with and consumed some walleye size classes. Walleye 14–30 cm and > 30 cm had higher δ^15^N than 8–13 cm walleye, yellow perch (5.0–22.0 cm), white sucker (*Catostomus commersonii*; 12.1–41.3 cm), cisco (13.2–16.9 cm), mimic shiner (4.6–6.0 cm), golden shiner (5.5–7.4 cm), spottail shiner (5.9–10.0 cm), and logperch (*Percina caprodes*; 4.6-7.4 cm) (pairwise, F = 18.1-18.4, p < 0.01).

Fish continued to compose most of the diet found in stomachs. Fish constituted 44% of diet in the 14–30 cm size class (including 6% yellow perch and 6% logperch, n = 13) and the whole diet of > 30 cm fish (n = 3, Figure 6). Young walleye (8–13 cm) had a different diet, and their δ^13^C values were close to those of benthic communities, mainly adult coleoptera (pairwise, F = 1.1, p = 0.3) and dipteran larvae (chironomids; pairwise, F = 3.5, p = 0.06). However, stomachs of young-of-the-year walleye predominantly contained fish (77% in total, including 18% logperch and 5% yellow perch, n = 20). Mayfly larvae, *Chaoborus* larvae, and crustaceans were present in 6–16% of stomachs sampled from 8– 13 cm and 14–30 cm walleye.

#### C2 (Vaudray Lake)

Unlike the other three lakes, the food web in C2 was not pyramid shaped (Figure 5d). Walleye > 30 cm competed for prey with northern pike (46.1–64.9 cm) and white sucker (35.3–54.7 cm) (δ^15^N and δ^13^C, pairwise, F = 0.5–0.8, p > 0.4). Walleye 14–30 cm long and trout-perch (6.0–9.3 cm) occupied the second to top trophic level. However, these species engaged in niche separation, with walleye being a pelagic predator and trout-perch being associated with macroinvertebrates and spottail shiner (2.6–9.0 cm) (pairwise, F (δ^15^N) = 1.6, p > 0.2, F (δ^13^C) = 11.3, p < 0.01). Yellow perch (5.8–24.2 cm) was the major fish prey for all length classes of walleye, northern pike and white sucker (pairwise, F (δ^15^N) = 25.6–49.7, p < 0.01; F (δ^13^C) = 0.8–2.2, p > 0.1). Large walleye (> 30cm), northern pike and white sucker also preyed on logperch (4.4–5.4 cm) (F (δ^15^N) = 41.1–184.8, p < 0.01; F (δ^13^C) = 0.7–2.3, p >= 0.1).

As in C1 but by contrast with the degraded lakes, the diet of young walleye (8–13 cm) in C2 differed from that of walleye > 30 cm long (pairwise, δ^13^C F = 4.4, p < 0.05). Young walleye also had higher δ^15^N values (pairwise, F = 7.1–261, p < 0.05) but similar δ^13^C values (pairwise, F = 1.9–2.7, p > 0.06) to all sizes of zooplankton. This result was consistent with young-walleye stomach contents, where *Mysis* sp. (pairwise, F = 0.78, p = 0.50) and *Chaoborus* larvae (pairwise, F = 1.75, p = 0.14) contributed to 4-30% of diets (n = 22, Figure 6). These macroinvertebrates were also found in the stomachs of walleye 14–30 cm (33% *Mysis* and 20% *Chaoborus,* n = 8) and > 30 cm (0% *Mysis* and 17% *Chaoborus*, n = 11). Diet was composed of 61% digested fish in stomachs from 8–13 cm walleye, 20% in stomachs of 14–30 cm fish, and 50% in stomachs of > 30 cm fish.

## Discussion

Our study demonstrated that walleye populations can be naturally sustained in the four lakes studied without the support of continuous stocking. However, degraded and undegraded lakes displayed different population dynamics.

Contrary to our hypothesis, the growth and relative abundance of young-of-the-year were higher in the degraded lakes than in control lakes. High early growth rates and greater relative abundance were associated with a greater abundance of daphniids, whose abundance is a determining factor in the growth and survival of walleye larvae (McDonnell and Roth 2014). The abundance of zooplankton in the degraded lakes of this study, particularly daphniids, is not supported by similar studies. For example, in the boreal Middle Lake (Sudbury, Canada), the abundance of cladocerans was limited in lakes polluted by metals (Yan et al. 2004). Our study showed that cladoceran abundance peaked during the spring in degraded lakes: but subsequently declined (Bosminidae and Holopedidae (*Holopedium gibberum*)) or were completely absent (Daphniidae in D1) by mid-summer (Gabriele et al. In prep; Figure S1). We hypothesized that Daphniids are in their resting egg life-stage during the summer when water quality deteriorates (Ringot et al. 2018). Trace-element concentrations in water increase from spring to summer (Manasypov et al. 2015), and zooplankton tend to have higher metal concentration in summer than in spring (Yan et al. 1989). This dynamic is advantageous for walleye larvae, which require abundant zooplankton in spring. The high spring abundance of *Daphnia* could compensate for the low biomass of calanoid copepods in degraded lakes compared to undegraded controls. Calanoids accumulate molecules that are important for fish health and growth (e.g., omega-3 and pigments) (Grosbois et al. 2017; Schneider et al. 2017), and their absence can produce nutrient deficits in juvenile fish (Mejri et al. 2021).

The abundant food resources for larvae in degraded lakes cannot fully explain high rates of growth and survival of young walleye until fall. The rapid growth of walleye in degraded lakes allows them to rise rapidly through the trophic levels. The swift growth of walleye in degraded lakes may reflect a strategy by which young-of-the-year transition rapidly from a zooplankton-based diet to piscivory, thereby avoiding reliance on macroinvertebrates, which may be limited in those ecosystems (Kövecses et al. 2005).

Stable isotope δ^15^N values also demonstrated that walleye was the apex predator in degraded lakes. This could be explained by the absence or reduced presence of other predatory fishes: predation on young walleye was limited to walleye > 30 cm long and northern pike (in D2 Lake only). Smallmouth bass (*Micropterus dolomieu*) which can prey on walleye young-of-the-year (Olivencia et al. 2024; Van Zuiden and Sharma 2016), was notably absent in degraded lakes (but present in C1). The high relative abundances of young-of-the-year walleye in degraded lakes lead to high relative abundance of adult in D1, but not in D2. Walleye in both lakes reproduced younger and had shorter lifespans than in control lakes, reflecting a similar strategy but different results. Adult mortality is therefore likely to be explained by different processes in D1 and D2 lakes.

The high production of young walleye and premature adult mortality mirrored previous observations of yellow perch that were made in metal contaminated lakes, including D1 and D2 (Pyle et al. 2008; Couture and Pyle 2008). Populations of predatory fish in these lakes appear to exhibit a “live fast, die young” strategy in which adult mortality may be linked to food limitation, metal concentrations, recreational fishing, or other habitat-related factors.

In metal contaminated lakes (including D1 and D2), yellow perch bottlenecks at small sizes due to the absence or low abundance of macroinvertebrates, which serve as the transitional food resource between zooplankton and fish (Sherwood et al. 2002). In our study, yellow perch was the most frequently found prey in walleye stomachs from degraded lakes (Figure 6). Isotopic values also demonstrated a strong predator-prey relation between walleye and yellow perch (Figure 5 a and b). In D1, the hypothesis of population limitation by a shortage of large prey items needs further investigation. In this lake, only small yellow perch, brown bullhead or walleye themselves (cannibalism) are available to larger walleye. By contrast, walleyes in D2 have access to a more diverse selection of fish species and prey sizes. Walleye in D2 also demonstrated some flexibility in their feeding habits. Adult walleye measuring > 30 cm showed a slight shift towards pelagic communities (lower δ^13^C values), relying more on cisco and golden shiner than smaller walleye. Therefore, our results do not support the food resources hypothesis to explain adult walleye mortality in D2.

The accumulation of metals in the muscles or organs of fish could contribute to adult mortality, as individuals need to allocate more energy towards detoxification (Sherwood et al. 2000). This diversion of energy could potentially hinder other essential functions, including growth, reproduction, and survival. Further studies are needed to understand the effects of metals on adult walleye, as high metal concentrations are present in D1 and D2 (Laflamme et al. 2000).

The photic zone was deeper in the degraded lakes of this study. Degraded lakes are often located in urban areas, and urbanization within their watersheds reduces the inputs of clay particles and tannins, which are known to naturally increase water turbidity (Grosbois et al. 2023, Hasan et al. 2023). Intensified human presence also makes these lakes more vulnerable to the introduction of invasive plants; *Myriophyllum spicatum* have been introduced in D1 and D2 (MELCCFP 2024b). Aquatic submerged macrophytes compete with phytoplankton and therefore reduce water turbidity (Thi Nguyen et al. 2015). D1, being shallow with a maximum depth of 8 m, provides favorable conditions for the growth and expansion of aquatic plants. Greater water clarity reduces the optimal light-temperature habitat for walleye, thereby reducing the population size of walleye and consequently affecting the harvest biomass of a sustainable fisheries (Lester et al. 2004; Hansen et al. 2019).

As walleye age, natural mortality decreases while fishing-related mortality increases (Hansen et al. 2011). Using data from experimental net inventories, it was possible to calculate total mortality (Mainguy and Moral 2021) and natural mortality using the methods described by Pauly (1980) and Jensen (1996), respectively, to estimate the mortality attributable to fishing in the four different study lakes (total mortality – natural mortality = fishing mortality) (Table S6).

Total mortality was higher in degraded Osisko (D1) (0.56) and Dufault (D2) (0.60) lakes than control Dufay (C1) (0.38) and Vaudray (C2) (0.30) lakes. Natural mortality accounted for the majority (34-35 %) of mortality in D1, while recreational fishing could be the primary cause of mortality for adult walleye in D2 (35–39% of mortality). D2 is situated in the city of Rouyn-Noranda and is easily accessible to anglers. Fishing mortality is estimated to be lower for the other three lakes, including D1 (16–21%) (Table S6). Although D1 is in downtown Rouyn-Noranda, motorized boats are not recommended, which reduces the vulnerability of walleye to recreational fishing. Additionally, its contamination with metals has given it a poor reputation and influences anglers’ decisions to release or retain specimens for personal consumption.

## Conclusion

Our study revealed important differences in the life cycle of walleye in degraded lakes compared to reference lakes. Comparable to yellow perch in the same lake (Couture and Pyle 2008), walleye have adopted a “live fast, die young” strategy: young-of-the-year are highly abundant, exhibit fast growth at an early age, and walleye reproduce at a younger age. This rapid start is supported by a high spring abundance of zooplankton, particularly cladocereans, a crucial food resource for larvae. Despite this surprising success in early life stages, the age structure of the populations reveals an underlying limitation: adults are, on average, younger in degraded lakes, and individuals older than 10 years are very rare, indicating that walleye live shorter lives than in reference lakes.

The causes of this mortality likely differ between the two degraded lakes. In D1, natural causes appear to dominate, particularly the limited availability of suitable food resources for large walleye. Indeed, yellow perch is the only available prey, but they remain small due to the lack of macroinvertebrates (Sherwood et al. 2002). In D2, recreational fishing is the most probable cause, according to our mortality calculations. The lake’s accessibility makes it more vulnerable to recreational fishing (Kaemingk et al. 2020), and fishing mortality data support this hypothesis.

This study demonstrated that stocking walleye can successfully restore a population. However, it also highlights that walleye self-sufficiency is compromised by a short adult stage in these ecosystems. This information will help guide management decisions for future interventions and/or studies in these lakes.

## Supporting information

supplentary-material-Blaney

## Acknowledgements

We would like to thank Alexane Gaudet and Marc-Olivier Roberge from MELCCFP for their valuable advice throughout the project. We extend our gratitude to the interns and students from GREMA who participated in field collections and laboratory work: Ariane Barrette, Lehann Bouchard, Julianne Breton, Olivier Bruneau, Yagmur Cakir, Marilou Cournoyer, Benjamin Ferron, Justin Gagnon, Marta Gabriele, Javier Gimenez Castillo, Mylène Gosselin, Éléa Jaskolski, Liv Jessen, Félix Labbé, Jade Lessard, Julie Marchal, Jérémy Mainville-Gamache, Chloé Tanguay, Antoine Villeneuve, and William Vincent. We are also grateful to volunteers Krystyn Chamberland, Alain Fort, and Guy Larochelle, who greatly assisted us with monitoring ice-breakup and spring temperatures in D1, D2 and C2. We thank Miguel Montoro-Girona for lending his drone and truck for the project. Additionally, we thank Olaloudé Judicaël Franck Ossé for assistance with statistical analyses. We also thank Technosub, OBVT, Collectif Territoire, and the Department of Fisheries and Oceans (DFO) for their assistance throughout the project. This project was supported by Hécla Quebec, NSERC-FCI Smart Forest, FRQNT, the FEDECP, and the Fondation Héritage faune. This study contributes to scientific programs of *Groupe de recherche interuniversitaire en Limnologie* (GRIL) and *Ressources aquatiques Québec* (RAQ).

## Notes

### Competing Interest Statement

The authors have declared no competing interest.

