## Supplementary material for "Live fast and die young: walleye populations adapt their life cycle to degraded lakes of the Canadian clay belt": supplentary-material-Blaney

Table S1: Walleye stocking made by MELCCFP in Osisko Lake

| **Date** | **Number** | **Stage** | **Lake source** |
| --- | --- | --- | --- |
| 1999-10-14 | 5 000 | Young-of-the-year (fingerling) | Unknown |
| 2000-10-25 | 4 130 | Young-of-the-year (fingerling) | Aylmer |
| 2001-10-17 | 1 122 | Young-of-the-year (fingerling) | Preissac |
| 2011-09-30 | 5 000 | Young-of-the-year (fingerling) | Dufay or Dasserat |
| 2012-10-16 | 2 000 | Young-of-the-year (fingerling) | Dufay |
| 2013-10-08 | 16 575 | Young-of-the-year (fingerling) | 3 000 from Dufay et 13 575 from Dasserat |
| 2014-10-07 | 5 000 | Young-of-the-year (fingerling) | Dufay |
| 2015-10-06 | 10 000 | Young-of-the-year (fingerling) | Dasserat |
| 2016-09-30 | 14 296 | Young-of-the-year (fingerling) | Dufay |
| 2017-09-21 | 11 080 | Young-of-the-year (fingerling) | Dufay |
| 2018-09-18 | 19 891 | Young-of-the-year (fingerling) | 7 610 from Dufay et 12 281 from Dasserat |

*Source : adapted from MELCCFP 2022*

Table S2: Walleye stocking made by MELCCFP in Dufault Lake

| **Date** | **Number** | **Stage** | **Lake source** |
| --- | --- | --- | --- |
| 1986 | 1 400 000 | Young-of-the-year (fry) | Réservoir Kipawa |
|  |  |  | Rivière Waswanipi |
|  |  |  | Rivière Rémigny |
| 1986 | 350 | Adult | Réservoir Kipawa |
|  |  |  | Rivière Waswanipi |
|  |  |  | Rivière Rémigny |
| 1987 | 60 | Adult | Duparquet |
| 2001 | 1 122 | Young-of-the-year (fingerling) | Preissac |
| 2008 | 4 521 | Young-of-the-year (fingerling) | Dufay |
| 2009 | 1 658 | Young-of-the-year (fingerling) | Dufay |
| 2010 | 7 511 | Young-of-the-year (fingerling) | Dufay |
| 2011 | 28 980 | Young-of-the-year (fingerling) | 26 700 from Dasserat and 2 280 from Dufay |
| 2012 | 14 495 | Young-of-the-year (fingerling) | Dufay |
| 2013 | 49 421 | Young-of-the-year (fingerling) | 28 921 from Dasserat et 20 500 from Dufay |
| 2014 | 15 000 | Young-of-the-year (fingerling) | Dufay |
| 2015 | 32 834 | Young-of-the-year (fingerling) | 1 580 from Dufay and 31254 from Dasserat |

*Source : adapted from MELCCFP 2022*

Table S3: Differentiating young-of-the-year walleye by length class during fieldwork (September 2023). Data shown for Age 0 are only for electrofishing and data for Age 1 also include gillnets (age 1 walleye were not captured during electrofishing).

|  | | Age | **Total length (cm)** | | | |
| --- | --- | --- | --- | --- | --- | --- |
|  |  |  | Mean | Standard error | Minimum | Maximum |
| Study lakes | Osisko | 0 (n = 34) | 14.5 | 1.1 | 12.5 | 16.5 |
|  |  | 1 (n = 32) | 24.0 | 2.7 | 20.1 | 32.0 |
|  | Dufault | 0 (n = 33) | 12.4 | 1.3 | 9.7 | 14.8 |
|  |  | 1 (n = 47) | 21.6 | 2.1 | 18.8 | 31.1 |
|  | Dufay | 0 (n = 28) | 10.7 | 0.8 | 8.6 | 12.4 |
|  |  | 1 (n = 15) | 16.2 | 2.4 | 20.2 | 12.8 |
|  | Vaudray | 0 (n = 21) | 9.1 | 0.9 | 7.5 | 10.9 |
|  |  | 1 (n = 27) | 14.7 | 1.8 | 19.9 | 12.3 |

Table S4: Number of species per lake, mean and range of total length of captured individuals per species.

| **Study lakes** | **Fish species (n)** | **Total Length (cm)**  mean (min – max) |
| --- | --- | --- |
| Osisko (D1) | Walleye (Age 0, n = 34) | 14.5 (12.5 – 16.5) |
|  | Yellow perch (n = 10) | 5.3 (4.0 – 6.9) |
| Dufault (D2) | Walleye (Age 0, n = 33) | 12.4 (9.7 – 14.8) |
|  | Golden shiner (n = 10) | 4.4 (3.1 – 5.5) |
|  | Logperch (n = 10) | 6.5 (6.1 – 8.6) |
|  | Trout-perch (n=10) | 7.4 (5.2 – 10.2) |
|  | White sucker (n = 10) | 7.2 (5.9 – 8.3) |
|  | Yellow perch (n = 11) | 7.1 (6.1 – 10.7) |
| Dufay (C1) | Walleye (Age 0, n = 28) | 10.7 (8.6 – 12.4) |
|  | Logperch (n = 8) | 6.5 (4.6 – 8.0) |
|  | Mimic shiner (n = 10) | 5.3 (4.6 – 6.4) |
|  | Spottail shiner (n = 10) | 9.1 (7.3 – 10.0) |
|  | Trout-perch (n = 10) | 7.7 (6.5 – 9.0) |
|  | Yellow perch (n = 10) | 5.6 (5.0 – 5.9) |
| Vaudray (C2) | Walleye (Age 0, n = 21) | 9.1 (7.5 – 10.9) |
|  | Logperch (n = 10) | 4.8 (4.4 – 5.4) |
|  | Spottail shiner (n = 10) | 5.6 (2.6 – 9.0) |
|  | Trout-perch (n = 10) | 7.6 (6.0 – 9.3) |
|  | Yellow perch (n = 10) | 6.3 (5.8 – 7.2) |

Table S5: Full result for A50 in each lake. “Not provided” means the GLM converged with the link family, but the McCullagh and Nelder test did not provide a p-value. “Did not converge” means the GLM did not converge.

| Lake | Sex | n | Link | McCullagh and Nelder GOF (p-value) | Osius and Rojek GOF (p-value) | AIC | A50 (from selected model) | 95% IC  (lwr : upr) | A50 decision (with model and maturation ogive) |
| --- | --- | --- | --- | --- | --- | --- | --- | --- | --- |
| Osisko | F | 101 | Logit | 0.01 | 0.984 | 51.72 |  |  |  |
|  |  |  | Probit | 0.007 | 0.999 | 54.75 |  |  |  |
|  |  |  | Cloglog | Not provided | 0.999 | 47.45 | 4.43 | 4 : 5 | **5** |
|  | M | 65 | Logit | Not provided | 0.999 | 7.82 | 2.03 | 2 : 2 | **3** |
|  |  |  | Probit | Not provided | 0.999 | 7.82 |  |  |  |
|  |  |  | Cloglog | Did not converge |  |  |  |  |  |
| Dufault | F | 109 | Logit | 0.5 | 0.998 | 16.21 |  |  |  |
|  |  |  | Probit | 0.5 | 0.999 | 16.68 |  |  |  |
|  |  |  | Cloglog | 0.5 | 0.999 | 15.43 | 5.52 | 5 : 7 | **6** |
|  | M | 133 | Logit | 0.06 | 0.794 | 77.81 |  |  |  |
|  |  |  | Probit | 0.05 | 0.976 | 77.32 | 4.10 | 4 : 5 | **4** |
|  |  |  | Cloglog | 0.005 | 0.999 | 85.20 |  |  |  |
| Dufay | F | 72 | Logit | 0.99 | 0.644 | 47.40 |  |  |  |
|  |  |  | Probit | 0.9 | 0.873 | 46.92 |  |  |  |
|  |  |  | Cloglog | 0.9 | 0.999 | 46.78 | 9.28 | 9 : 11 | **9** |
|  | M | 66 | Logit | 0.4 | 0.756 | 50.96 |  |  |  |
|  |  |  | Probit | 0.4 | 0.936 | 50.61 | 5.82 | 5 : 7 | **6** |
|  |  |  | Cloglog | 0.2 | 0.999 | 53.00 |  |  |  |
| Vaudray | F | 104 | Logit | 0.2 | 0.929 | 27.85 |  |  |  |
|  |  |  | Probit | 0.06 | 0.999 | 29.93 |  |  |  |
|  |  |  | Cloglog | 0.4 | 0.999 | 26.86 | 8.72 | 8 : 10 | **9** |
|  | M | 138 | Logit | 0.005 | 0.522 | 97.46 | 5.61 | 5 : 6 | **6** |
|  |  |  | Probit | 0.0007 | 0.992 | 101.75 |  |  |  |
|  |  |  | Cloglog | Did not converge |  |  |  |  |  |

Figure S1: Seasonal variations (spring vs summer) in zooplankton densities. The general trend is that different families of cladocerans are more abundant in degraded lakes during spring, and they decline over the summer (the opposite is observed in control lakes). No Daphniidae were found in the summer samples from Lake Osisko. Cyclopidae dominate in degraded lakes, while Calanoida dominate in control lakes.

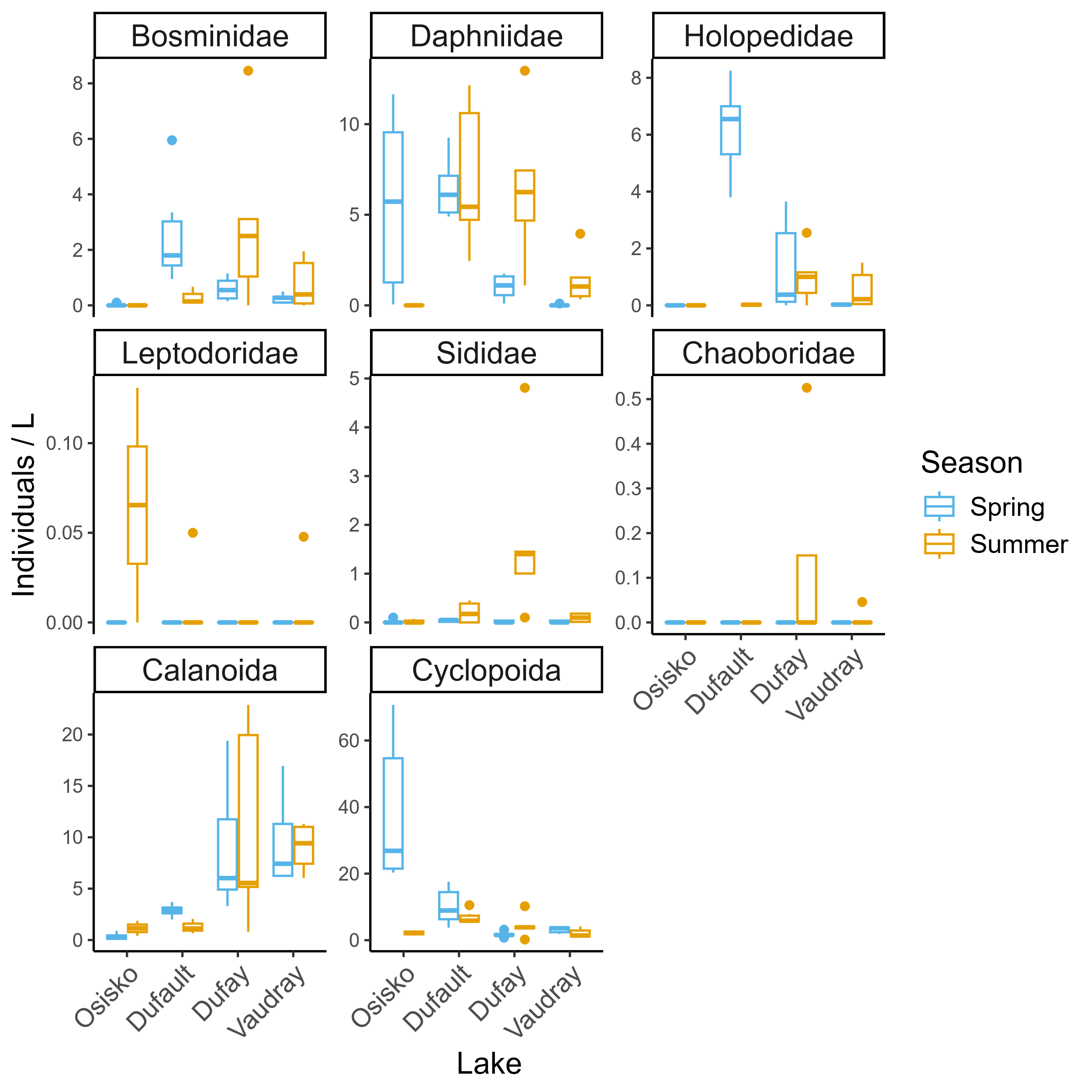

Figure S2: Biomass differences for Daphniidae in all 4 lakes and station. Degraded lakes have higher Daphniidae biomass, except one station in Osisko Lake

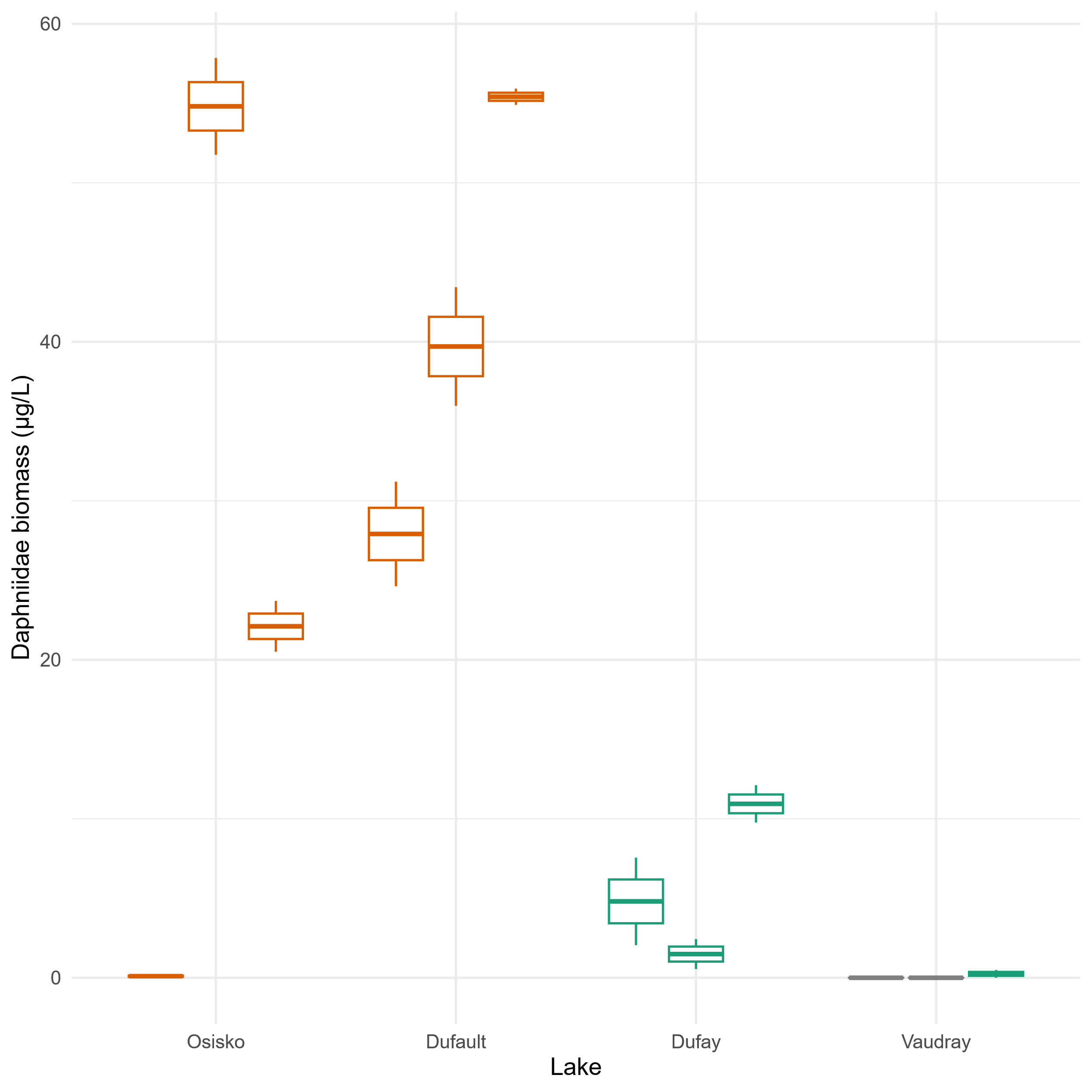

Table S6: Fishing mortality

| Lake | Total mortality (Z_obs_) | Natural mortality  (M)  Pauly 1980 / Jensen 1996 | Fishing Mortality  (Z_obs_ – M) |
| --- | --- | --- | --- |
| Osisko | 0.56 | 0.35 / 0.35 | 0.16 - 0.21 |
| Dufault | 0.60 | 0.22 / 0.18 | 0.35 – 0.39 |
| Dufay | 0.38 | 0.22 / 0.17 | 0.13 – 0.21 |
| Vaudray | 0.30 | 0.22 / 0.17 | 0.08 – 0.14 |

Total mortality (MainGuy et Moral 2021)

There isn’t a specific equation as such; it’s more of a process. The detailed process is explained in MainGuy and Moral in the appendices. A selection of models was performed among the distribution families that showed good convergence (using the hnp test presented in MainGuy and Moral (2021)). The Zobs value was calculated by performing model averaging using the weights from the model selection table.

Natural mortality

*Pauly method (1980)*

$$M= e^{-0.0152+0.654\ln k-0.279\ln L_{\infty}+0.4634\ln T^{o}}$$

Where M is natural mortality, k is the growth parameter from the Von Bertalanffy equation. L_∞_ is the asymptotic length from Von Bertalanffy equation and T^o^ is the annual average temperature.

It is recommended to use an annual temperature mean of 10 °C for walleye inland lakes (Dianel Nadeau, MRNF, Rouyn-Noranda, Canada, personal communication).

*Jensen method (1996)*

M = 1,5k

Where k is the parameter of growth from the Von Bertalanffy equation.

Fishing mortality

Fishing mortality is estimated as the difference between total mortality (Z_obs_) and natural mortality

$${Fishing}_{mortality}= Z_{obs}-M$$
